# A Biophysical Model of Human Colonic Motor Pattern Generation in Health and Disease

**DOI:** 10.64898/2026.04.15.718795

**Authors:** Ahilan Anantha Krishnan, Phil G. Dinning, Maria A. Holland

**Affiliations:** Department of Aerospace and Mechanical Engineering, University of Notre Dame, Notre Dame, 46556, IN, USA; Department of Surgery, Flinders Medical Centre, Bedford Park, 5042, SA, Australia; Bioengineering Graduate Program, University of Notre Dame, Notre Dame, 46556, IN, USA

**Keywords:** Colon motility, Manometry, diarrhea-predominant irritable bowel syndrome, slow-transit constipation, biophysical modeling

## Abstract

**Purpose:** Colonic motility disorders, including diarrhea-predominant irritable bowel syndrome and slow-transit constipation, impose a major clinical burden. Although high-resolution colonic manometry reveals characteristic spatiotemporal motor patterns, such as high-amplitude propagating contractions and cyclic motor pattern in healthy individuals, these patterns are often altered or absent in disease. Understanding how these patterns arise from underlying pacemaker, neural, and mechanical mechanisms is essential for improving treatment strategies.

**Methods:** We developed a biophysical whole-colon model that integrates an Interstitial Cells of Cajal–inspired oscillator network, enteric nervous system reflexes, a pressure-gated modulation element motivated by rectosigmoid brake behavior, and a nonlinear tube law describing colon wall mechanics. The model simulates spatiotemporal pressure patterns along the colon and allows systematic variation of physiological parameters associated with pacemaker activity, neural reflex control, and distal gating.

**Results:** A small set of parameters reproduces three illustrative motility patterns corresponding to healthy motility, diarrhea-predominant irritable bowel syndrome, and slow-transit constipation. The simulated pressure maps recapitulate key features observed in high-resolution manometry, including propagation direction, regional patterning of contractions, and case-specific changes in amplitude and coordination. Sensitivity analysis suggests that proximal excitation strength and waveform morphology strongly influence global motility metrics.

**Conclusion:** Our study presents a simple, biophysical framework for reproducing clinically observed colonic motor patterns and exploring their disruption in disease. More broadly, the model may help interpret clinical manometry in mechanistic terms and support hypothesis-driven *in silico* studies of colonic motility disorders.

## 1 Introduction

Colonic motility disorders, such as irritable bowel syndrome (IBS) and chronic constipation, impose a major clinical and societal burden. IBS affects roughly 10% of the global population and is a leading cause of outpatient visits, lost productivity, and reduced quality of life [1, 2]. Chronic constipation, encompassing both functional and idiopathic forms, affects an estimated 14-16% of adults in North America and Europe, particularly older adults and women [3, 4]. Chronic constipation contributes substantially to healthcare costs through both direct expenses (e.g., frequent healthcare use and medication) and indirect costs (e.g., reduced work productivity and functional impairment) [5, 6]. Despite their prevalence and impact, the mechanisms underlying these motility disorders remain incompletely understood, complicating diagnosis and treatment.

The healthy colon exhibits a diverse set of motor behaviors coordinated to ensure effective mixing, fluid absorption, and the eventual transit of fecal contents. High-amplitude propagating contractions (HAPCs) are episodic, high-pressure waves that propagate over long colonic distances and mediate mass movements, often triggering defecation [7–9]. In contrast, cyclic propagating motor pattern (CMP) occur more frequently in the distal colon, exhibit predominantly retrograde propagation, and contribute to content mixing and continence in a manner consistent with the proposed ‘rectosigmoid brake’ described in high resolution manometry studies [9–14]. These patterns are characterized by specific frequencies and directionalities, and their regulation reflects a complex interplay between the interstitial cells of Cajal (ICC), smooth muscle, and neural inputs. High-resolution manometry has enhanced the ability to observe these patterns *in vivo*, revealing nuanced spatial and temporal features that underlie physiological and pathological motility [9, 15].

The aforementioned dominant motor patterns observed in healthy individuals are reduced or altered in patients with IBS-D and slow-transit constipation (STC) [16–18]. For instance, Wiklendt et al. [16] report that IBS-D patients exhibit inhibited postprandial retrograde cyclic motor pattern activity in the distal colon, with a loss of the normal retrograde dominance seen in healthy subjects. This attenuation of retrograde propagation was hypothesized to permit premature rectal filling and contribute to urgency and diarrhea. In contrast, high-resolution colonic manometry in slow-transit constipation demonstrates an absence of the normal postprandial increase in cyclic motor patterns observed in healthy subjects [17].

These findings highlight the spectrum of motor abnormalities, beneath which lie a complex interplay of neuromuscular and sensory dysfunctions. For instance, IBS-D involves not only altered motility but also visceral hypersensitivity and altered reflex responses, implicating central and peripheral neural pathways [19–21]. Meanwhile, STC is considered to have subtypes with enteric neuropathy, myopathy, or depletion of the interstitial cells of Cajal, each associated with distinct patterns of hypomotility and therapeutic responsiveness [22–24]. Thus, understanding how pacemaker, neural, and mechanical mechanisms interact to generate colonic motor patterns is essential for improving treatment options. However, isolating their individual contributions in vivo remains challenging, as clinical manometric measurements reflect the combined output of overlapping sensory, neuromotor, and structural inputs, with substantial inter-individual variability [25]. This coupling complexity motivates the need for systems-level frameworks that can translate the motor pattern trends into mechanistically informed, patient-specific therapeutic strategies.

Computational modeling is increasingly seen as a valuable complement to experimental studies. Mechanistic *in silico* models can integrate known physiology to simulate colonic motor patterns across space and time, enabling the controlled exploration of hypotheses that are impractical to test experimentally [26, 27]. Such models could potentially serve as virtual platforms to investigate how varying physiological parameters influence emergent motor patterns, providing insights beyond the reach of current *in vivo* experiments. Previous modeling approaches have captured key aspects of colonic motility. Oscillator chain models inspired by the ICC network have demonstrated how frequency gradients and coupling influence wave coherence and propagation [28, 29]. Neurogenic models have emphasized enteric reflex control, simulating peristalsis via ascending excitation and descending inhibition [26, 30]. While informative, many of these models overlook critical features such as regional specialization (proximal vs. distal function), reflex asymmetry (oral excitation and anal inhibition), and the integration of mechanosensory feedback. As a result, they often fail to reproduce the characteristic high-resolution manometry patterns, such as HAPCs, CMPs, and rectosigmoid brake type dynamics observed in humans. Additionally, all existing modeling efforts have been restricted to short segments or based on animal physiology, limiting their translational relevance to human whole-colon motor patterns.

Here, we present a biophysical model of the human colon that integrates ICC-inspired oscillator dynamics, enteric reflexes, and pressure-mediated gating of distal activity to simulate clinically observed motor patterns. The colon is partitioned into proximal and distal regions, each with distinct pacing and reflex architectures. To capture the brake-like behavior reported in high-resolution manometry, we include a phenomenological gating element that modulates distal activity, enabling the model to reproduce key features such as anterograde HAPCs, retrograde CMPs, and the disruptions of these patterns observed in IBS-D and STC. By allowing adjustable parameters, the model supports hypothesis testing and offers insights into the coordination of colonic motility.

## 2 Materials and Methods

An overview of example high-resolution manometry recordings and the corresponding computational model architecture is shown in Fig. 1. We model the human colon as a one–dimensional chain of *N* = 75 segments, representing a total length of *L* =150 cm (2 cm per segment), corresponding to the average adult colon length [31]. The chain is partitioned into a proximal region (*N*_HAPC_ = 50) parameterized to reproduce HAPC-dominant long-range propagating events and a distal region (*N*_CMP_ = 25) parameterized to reproduce the cyclic propagating motor pattern (CMP) [7, 13, 32]. Anatomical landmarks are indexed along the chain: the cecum is at *s* = 0, the splenic flexure (SF) is at *s* = 50, and the rectosigmoid junction (RSJ) is at *s* = 75 (distal boundary). Motivated by distal brake-like behavior reported in high resolution manometry studies, we include a pressure sensing field (Fig. 1) to enable modulation of the distal activity [12–14]. Each segment *s* carries a phase *ϕ*_*s*_(*t*) (oscillator), an excitatory drive *E*_*s*_(*t*) ∈ [0, 1], a filtered drive 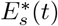, muscle activation *M*_*s*_(*t*) ∈ [0, 1], normalized cross–sectional area *A*_*s*_(*t*), and pressure *p*_*s*_(*t*).

**Fig. 1.**
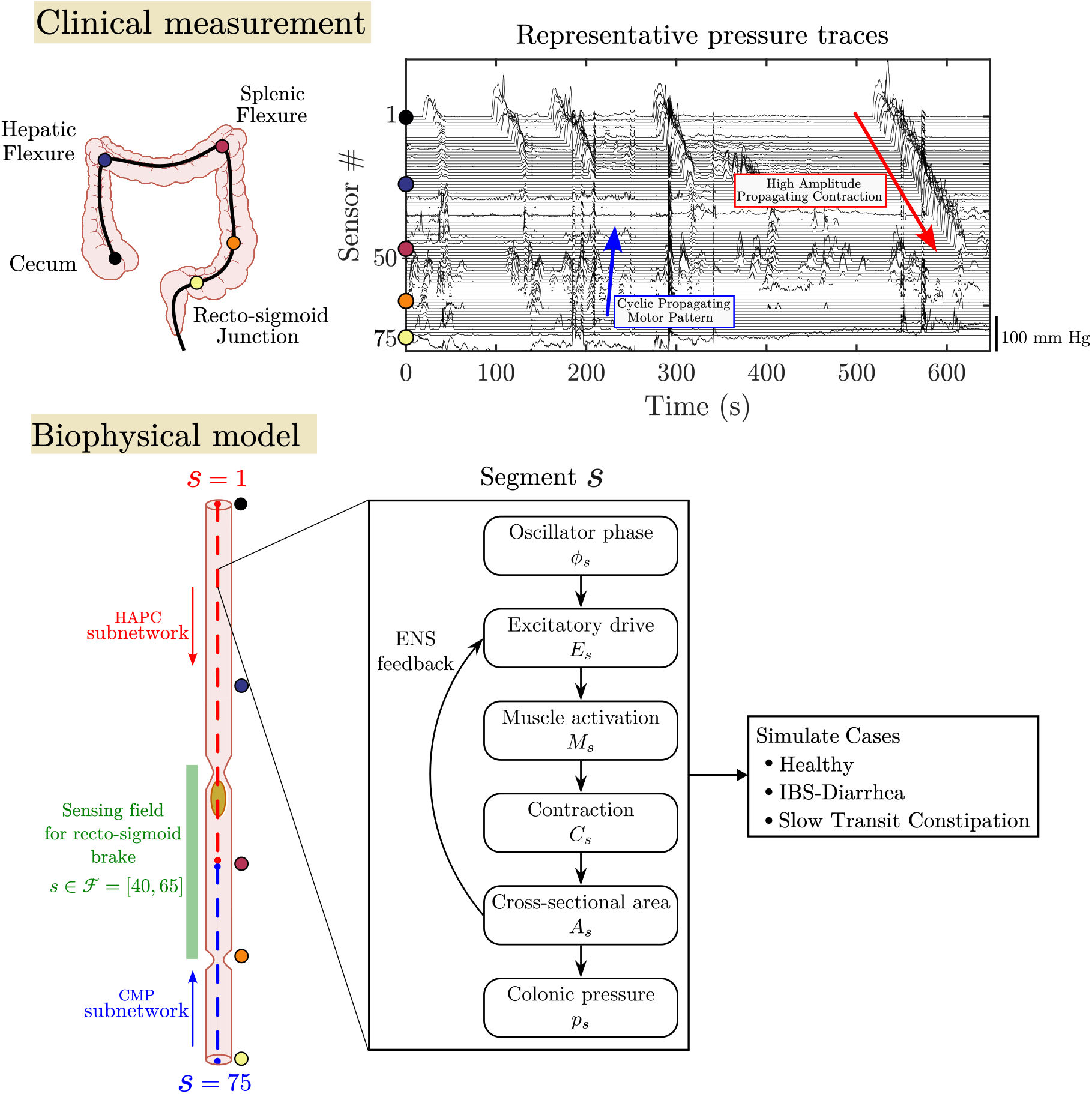
Overview of clinical and computational model for studying colonic motility (Top right) Clinical high-resolution manometry traces from a healthy subject show key motor patterns, including high-amplitude propagating contractions (HAPCs) and retrograde cyclic motor pattern (CMP) (Top left) Colored dots highlight anatomical locations along the colon from cecum to rectosigmoid junction and the corresponding locations on the manometry plot (Bottom) Schematic of the biophysical model architecture The colon is modeled as a chain of 75 segments, each containing a coupled oscillator (*ϕ*_*s*_), enteric reflex control (*E*_*s*_), muscle activation (*M*_*s*_), contraction (*C*_*s*_), and pressure state (*p*_*s*_) The model can simulate healthy, IBS-D-like, and STC-like motility patterns

### 2.1 Oscillator model of ICC dynamics

ICC, the gut’s pacemaker cells, generate rhythmic slow-wave depolarizations that underlie colonic motor patterns. These cells form electrically coupled networks that behave like chains of locally interacting, noise-influenced oscillators [15, 33]. This organization motivates our use of a nearest-neighbor, noise-sensitive oscillator chain to represent colonic pacemaker activity. In this framework, each colonic segment is modeled as a phase oscillator, and the colon is partitioned into two subnetworks with distinct physiological roles and propagation directions.

- **Proximal HAPC subnetwork:** Segments *s* = [1, 50] oscillate at a low intrinsic frequency *f*_HAPC_. Oscillators are coupled anterogradely, producing traveling HAPCs. In the adult colon, HAPCs are typically triggered episodically by reflexes (e.g., gastrocolic response) [32]; however, we approximate those events here with a continuous low-frequency oscillator for simplicity, representing a phenomenological propulsive pattern generator.
- **Distal CMP subnetwork:** Segments *s* = [51, 75] oscillate at a higher frequency *f*_CMP_, producing retrograde cyclic contractions consistent with the rectosigmoid brake behavior reported in the literature [12–14]. Segments are coupled retrogradely [13].

Our model builds on the Kuramoto model [34], which describes synchronization in oscillator networks. Unlike the standard Kuramoto model with global coupling, we implement nearest-neighbor interactions along a one-dimensional chain to capture the diffusive coupling observed in excitable media, such as colonic slow-wave conduction [35]. A stochastic phase term introduces intrinsic noise to the phase, which affects wave coherence and stability. The phase evolution for any colonic region *κ* ∈ {HAPC, CMP} is given by the generic form

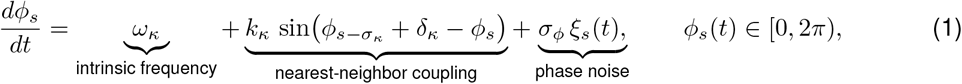

where *ω*_*κ*_ = 2*π*(*f*_*κ*_/60) is the intrinsic angular frequency (rad/s), with *f*_*κ*_ specified in cycles per minute (cpm), *k*_*κ*_ is the coupling strength, δ_*κ*_ is the per-segment phase lag, and *ξ*_*s*_(*t*) is Gaussian white noise scaled by the phase-drift amplitude *σ*_*ϕ*_ (see Table 2). Propagation direction is prescribed by the neighbor index *s* − *σ*_*κ*_ in Eq. (1):

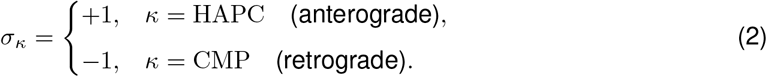

The phase lag δ_*κ*_ encodes the preferred propagation speed *v*_*κ*_:

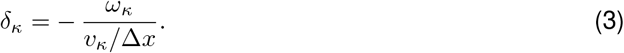

with the segment length Δ*x* = *L*/*N*. By tuning *f*_*κ*_, *k*_*κ*_, δ_*κ*_, and the direction selector *σ*_*κ*_, the model reproduces region-specific ICC dynamics: proximal segments exhibit lower-frequency, long-range propagating waves, while distal segments favor higher-frequency, retrograde activity. These distinctions reflect spatial heterogeneity in ICC networks, including variations in connectivity and intrinsic pacing, and give rise to heterogeneous slow-wave propagation across the colon [28, 33].

### 2.2 Pacemaker waveform

ICC generate slow-wave electrical rhythms in the human colon, characterized by a rapid depolarizing upstroke followed by a slower decay [36]. To capture this asymmetry, each oscillator converts its phase into a skewed waveform *W*_*s*_:

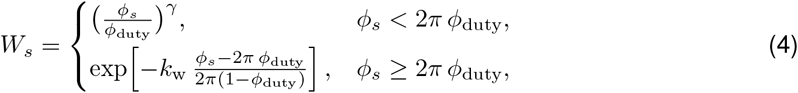

where *ϕ*_duty_ is the duty fraction, *γ* the upstroke sharpness, and *k*_w_ the decay constant. *W*_*s*_ represents the pacemaker output of the ICC, which generates slow-wave electrical rhythms at approximately 2–6 cycles per minute throughout the human colon (Table 1) [15].

**Table 1.** Global waveform, muscle-model parameters and non-linear tube-law constants for the whole-colon motility model. Parameter values were chosen to be physiologically plausible and consistent with reported ranges where available or selected phenomenologically to reproduce characteristic features of clinically measured pressure maps.

| Parameter | Description | Value [Source] |
| --- | --- | --- |
| $f_{\text{HAPC}}$ [cpm] | HAPC pacemaker frequency | 0.4 [25] |
| $f_{\text{CMP}}$ [cpm] | CMP pacemaker frequency | 3.0 [7] |
| $v_{\text{CMP}}$ [cm/s] | Target CMP propagation speed | 5.0 [7] |
| $\phi_{\text{duty}}$ [-] | Duty fraction of $2\pi$ | 0.1 |
| $\gamma$ [-] | Waveform upstroke sharpness | 2 |
| $k_w$ [-] | Exponential decay constant | 8 |
| $a_{\text{CMP}}$ [-] | CMP pacemaker amplitude | 0.75 |
| $\alpha_{\text{spread,HAPC}}$ [-] | Proximal spread coefficient | 0.1 |
| $\alpha_{\text{spread,CMP}}$ [-] | Distal spread coefficient | 0.05 |
| $\alpha_{\text{asc}}$ [-] | Ascending (oral) reflex gain | 0.1 |
| $\alpha_{\text{desc}}$ [-] | Descending (anal) reflex gain | 0.08 |
| $\tau_E$ [s] | Excitatory filter constant | 1.0 |
| $\tau_{\text{on}}$ [s] | Muscle contraction constant | 1.0 [46] |
| $\tau_{\text{off}}$ [s] | Muscle relaxation constant | 1.2 [46] |
| $k_c$ [-] | Contraction–curve steepness | 6.0 |
| $M_{50}$ [-] | Contraction–curve midpoint | 0.3 |
| $\beta_C$ [-] | Max. fractional area reduction | 0.8 |
| $p_{\text{ref}}$ [mmHg] | Reference pressure | 1 |
| $E_w$ [mmHg] | Wall stiffness | 25 |
| $n_p$ [-] | Tube law exponent | 1.3 |
| $p_{\text{on}}$ [mmHg] | Pressure threshold for brake ON | 50 |
| $p_{\text{off}}$ [mmHg] | Pressure threshold for brake OFF | 20 |
| $\tau_P$ [s] | Pressure filter constant | 1.2 |

**Table 2.** Case–specific model parameters for healthy, IBS-D, and STC.

| Parameter | Description | Healthy | IBS-D | STC |
| --- | --- | --- | --- | --- |
| $v_{\text{HAPC}}$ [cm/s] [7] | Preferred HAPC propagation speed | 1.2 | 5.0 | 1.2 |
| $k_{\text{HAPC}}$ [-] | Coupling strength (HAPC) | 2.0 | 2.0 | 0.2 |
| $k_{\text{CMP}}$ [-] | Coupling strength (CMP) | 2.0 | 2.0 | 0.2 |
| $a_{\text{HAPC}}$ [-] | HAPC pacemaker amplitude | 0.77 | 0.70 | 0.62 |
| $\sigma_\phi$ [rad/$\sqrt{\text{s}}$] | Phase noise amplitude | 0.0 | 0.0 | 0.5 |
| Brake status [7, 47] | State of rectosigmoid brake | Active | Disabled | Active |

### 2.3 Excitatory drive

The total excitatory drive in each segment is composed of two parts: the intrinsic ICC slow-wave (myogenic pacing) component and the contributions from enteric reflexes.

#### 2.3.1 ICC contribution

For segment *s* belonging to subnetwork *κ* ∈ {HAPC, CMP}, the ICC slow-wave waveform *W*_*s*_ is converted into regional slow-wave activation (myogenic pacing signal).

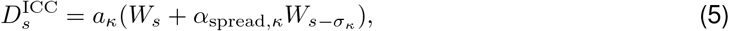

where *a*_*κ*_ scales the regional magnitude of the ICC-derived slow-wave input used in the model (Table 1). In the proximal region, a larger value of *a*_HAPC_ provides a stronger propulsive activation consistent with the higher pressures observed during HAPCs, while acknowledging that, *in vivo*, recruitment into high-amplitude HAPCs is strongly shaped by enteric excitation and mechanical context rather than by slow waves alone. The distal region uses a smaller value (*a*_CMP_), consistent with lower-amplitude cyclic activity. For each subnetwork, *α*_spread,*κ*_ is the local spread coefficient.

#### 2.3.2 Enteric nervous system (ENS) contribution

To represent the polarized peristaltic reflex, in which luminal distension evokes ascending excitation on the oral side and descending inhibition on the anal side, we use local changes in cross-sectional area as a proxy for mechanosensory input. We define the local distension signal at the interface between segments *s* and *s* + 1 as the area gradient (where *A*_*s*_ denotes the local cross-sectional area):

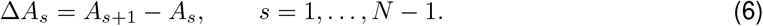

A positive gradient Δ*A*_*s*_ *>* 0 therefore indicates an increasing luminal area in the anal direction, consistent with the distension that triggers the polarized peristaltic reflex. Because each interface contributes to the two adjacent segments, the net ENS contribution to segment *s* is

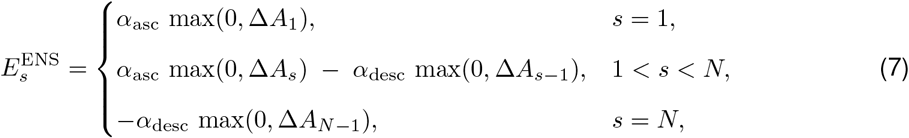

where *α*_asc_ and *α*_desc_ are the ascending (oral) and descending (anal) reflex gains, respectively (Table 1). Thus, our formulation implements the polarized peristaltic reflex (law of the intestine) [37, 38].

#### 2.3.3 Total excitatory drive

The effective excitatory drive in segment *s* combines ICC-derived slow-wave input and ENS reflex contributions, and is regionally gated by a binary gate (a brake-inspired phenomenological abstraction) Γ_*κ*_(*t*):

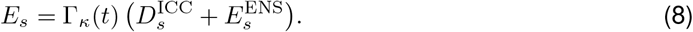

The proximal segments (Γ_HAPC_ = 1) always receive the full ICC and ENS inputs, whereas in the distal CMP region, the rectosigmoid brake can transiently switch off the entire excitatory drive when Γ_CMP_(*t*) = 0.

### 2.4 Muscle activation

Colonic smooth muscle exhibits faster active contraction than relaxation [39], and it does not respond instantaneously to rapid fluctuations in neural excitation. So, excitatory drive is low–pass filtered (smoothed) by

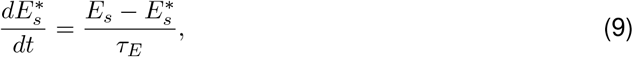

and then converted to muscle activation with asymmetric kinetics:

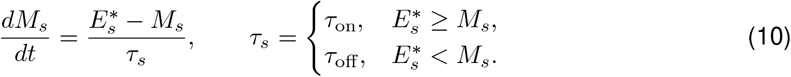

Together, this low-pass filtering of *E*_*s*_ to 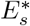 and the asymmetric time constants *τ*_on_ and *τ*_off_ capture the empirically faster contraction and slower relaxation of colonic smooth muscle [40] and provide a physiological smoothing of neural excitation, ensuring that the muscle response reflects the viscoelastic nature of the tissue (Table 1) [41].

### 2.5 Tube law

Muscle activation reduces the luminal area through the circumferential shortening of the wall, thereby elevating intraluminal pressure [42]. The contraction curve is defined as

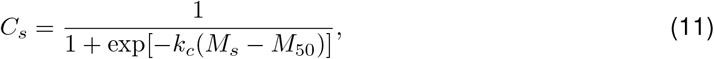

with steepness *k*_*c*_ and midpoint *M*_50_ (Table 1). The normalized cross-sectional area decreases as

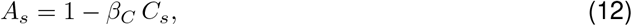

where *β*_*C*_ sets the maximum fractional reduction. Segmental pressure follows a nonlinear tube law that reproduces the steep stiffening observed at high distension [43]:

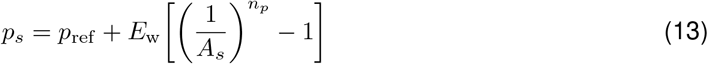

### 2.6 Rectosigmoid brake

In this model, distal CMP activity is modulated by a wide-field, pressure-dependent gate distributed over the left colon, representing the rectosigmoid brake behavior reported in high resolution manometry studies [12–14] (Fig. 1). Low–pass filtered pressure 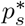 is first computed within the sensing field ℱ = [40, 65] as

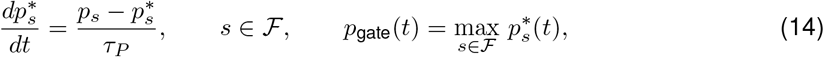

where *τ*_*P*_ is the pressure filter time constant, and *p*_gate_(*t*) represents the maximal filtered pressure in the sensing field. The rectosigmoid brake is modeled by a binary gate Γ_CMP_(*t*) ∈ {0, 1} that controls distal CMP activity. The gate closes (brake ON, Γ_CMP_ = 0) when the filtered pressure exceeds an activation threshold *p*_on_ and opens (brake OFF, Γ_CMP_ = 1) only after the pressure has fallen below a lower deactivation threshold *p*_off_. This is given by,

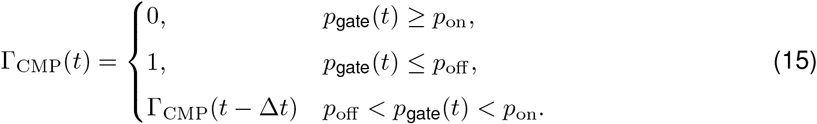

When the brake is ON (Γ_CMP_ = 0), distal excitatory drive is gated off by applying

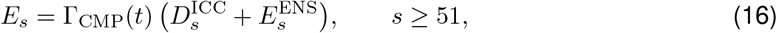

so that ICC- and ENS-derived inputs do not activate distal muscle during braking. The underlying distal oscillator phases continue to evolve, but their effect on excitation is suppressed by the gate.

### 2.7 Initial conditions

Simulations begin from a quiescent state. The proximal HAPC chain is initialized with a linear phase gradient chosen to produce a target propagation speed *v*_HAPC_, while the distal CMP chain is initialized with a uniform phase. The phase in segment *s* at *t* = 0 is

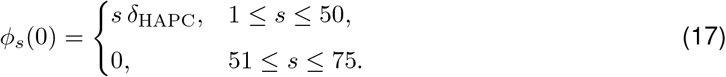

At *t* = 0, the remaining state variables are initialized consistently with the pacemaker output:

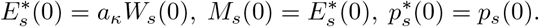

### 2.8 Simulating Illustrative Cases

The full model is implemented in Python 3.13. All cases use forward Euler integration with a time step of Δ*t* =0.05 s over a simulated duration of 150 s, starting from the above initial conditions. The stochastic term is discretized using the Euler-Maruyama method, so that phase increments scale with 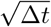 to ensure variance proportional to time [44]. In all simulations, consistent values were used for most parameters, as listed in Table 1. The cases differ by a select set of six parameters, with the full set of three case-specific values given in Table 2.

#### 2.8.1 Healthy

The healthy parameter set was chosen to represent coordinated colonic motility, supported by coupled pacemaker activity and intact enteric reflexes, along with moderate coupling to allow orderly propagation. The rectosigmoid brake was active and pressure-gated, suppressing distal activity when luminal pressure exceeded the designated threshold, thereby replicating the inhibitory modulation observed in a healthy distal colon [12]. No phase noise was added, so the pacemaker network operated in a stable, deterministic regime.

#### 2.8.2 IBS–D

To represent an IBS-D-like motility pattern, we substantially increased the HAPC propagation speed and slightly reduced the amplitude *a*_HAPC_ compared to the healthy case. The rectosigmoid brake was disabled (Γ_CMP_ ≡ 1), removing distal gating altogether and allowing continuous CMP activity. Reflex gains were unchanged, and no noise was added. These adjustments are intended to test whether the reduced brake-like distal gating can reproduce a pattern observed in some IBS-D patients [16].

#### 2.8.3 STC

The slow-transit constipation model preserved most of the baseline parameters but imposed very weak coupling and reduced proximal pacemaker amplitude, substantially limiting coordinated propagation. Phase noise was added to disrupt synchrony and introduce irregularity in the pacemaker rhythm. The rectosigmoid brake remained active, though the diminished upstream drive rendered its influence minimal. This parameter combination reflects the low-coherence, low-pressure motor patterns characteristic of clinically observed STC [25, 45].

## 3 Results

### 3.1 Simulated pressure maps

We simulated colonic motility across three illustrative cases corresponding to healthy motility, IBS-D, and STC (Figure 2, right). The model reproduces distinct region-specific motility patterns observed in high-resolution manometry.

**Fig. 2.**
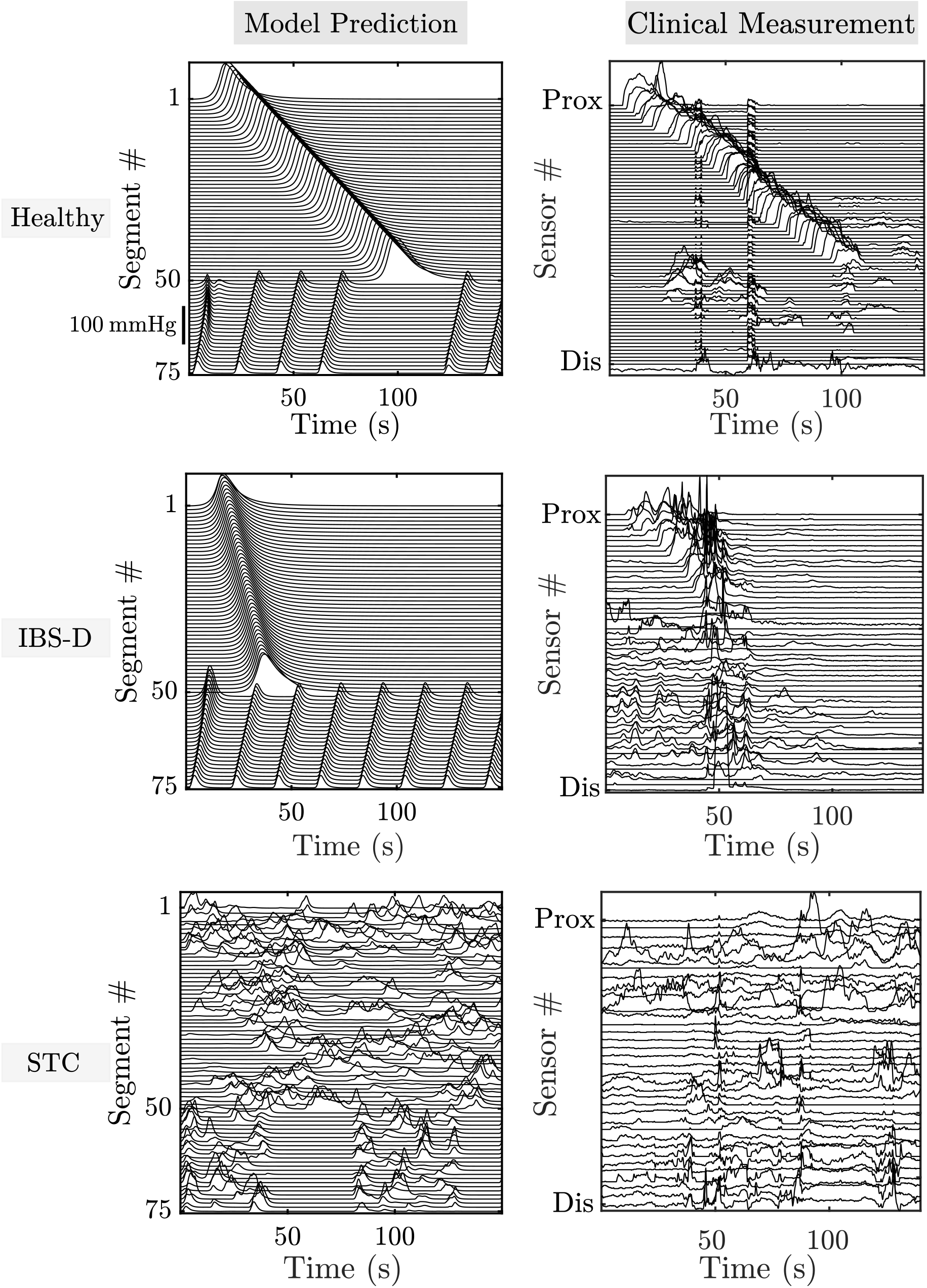
Clinically measured and simulated pressure maps across three illustrative cases Rows display illustrative examples corresponding to healthy motility (top), IBS-D (middle), and STC (bottom) Model-predicted pressure maps are shown on the left, with clinical measurements for corresponding examples on the right Sensors are indexed by anatomical position from proximal to distal colon; numbering reflects relative location because sensor counts used varied between subjects The model reproduces key features of regional motor patterns, including anterograde HAPCs, retrograde CMPs, and case-specific disruptions in coordination and amplitude

In the healthy case, coordinated anterograde HAPCs were reproduced in the proximal colon, and retrograde CMPs were reproduced in the distal colon. The rectosigmoid brake appropriately gates CMP activity based on mechanosensory feedback. The maximum simulated pressure was 142.4mmHg. In the illustrative IBS-D case, a frequent contraction with disrupted brake function was predicted with a peak pressure of 124.1mmHg. By contrast, the illustrative STC model showed impaired propagation and reduced coordination, modeled via increased phase noise to reflect disrupted pacemaker coherence and ICC dysfunction. Despite an intact brake, distal CMPs were irregular and fragmented. The peak pressure was markedly lower at 73.8mmHg, consistent with ineffective motility.

### 3.2 Comparison with clinical measurement

We compared the simulations with high-resolution manometry recordings from three example subjects obtained in our previous studies: a healthy male (26 years)[7], a female IBS-D patient (42 years) [16], and a constipated male child (10 years) [17]. Cropped segments were selected to align with the simulated time windows. As shown in Figure 2, the experimental traces display spatiotemporal patterns that the model successfully captures. In the healthy subject, pressure waves were well-organized and propagated distally, with peak pressures near 143.8mmHg. The IBS-D trace exhibited disorganized, frequent contractions and a peak of 125.3mmHg. The constipated subject showed fragmented, low-amplitude events with a maximum pressure of 74.0mmHg. The model thus reproduced key qualitative features of each illustrative case, including propagation direction, frequency, and regional dominance, supporting its physiological relevance. A quantitative comparison of maximum pressures between model predictions and clinical measurements is provided in Table 3.

**Table 3.** Maximum intraluminal pressures from model predictions and clinical measurements across cases. Values illustrate overall agreement between simulations and experimental data.

| Case | Model prediction (mmHg) | Clinical measurement (mmHg) |
| --- | --- | --- |
| Healthy | 142.4 | 143.8 |
| IBS-D | 124.1 | 125.3 |
| STC | 73.8 | 74.0 |

### 3.3 Sensitivity to model parameters

To examine the mechanisms and robustness, we perturbed model parameters and measured clinical manometric outcomes: (i) peak pressure (mmHg) and (ii) motility index, i.e., the area under the pressure–time curve (mmHg.s). The healthy case served as the baseline, and the sensitivity analysis was evaluated relative to this normal motility benchmark (Fig. 3). Note that parameters associated with the tube law were not included in the sensitivity analysis, as they primarily modulate the local pressure–area relationship and do not alter the spatiotemporal organization of propagating motor patterns. The model was most sensitive to the proximal wave amplitude (*a*_HAPC_), and the motility index was additionally sensitive to proximal pacemaker frequency (*f*_HAPC_) and waveform shape (decay constant *k*_w_). Secondary sensitivities were observed for proximal spread *α*_spread,HAPC_, ascending reflex gain *α*_asc_, and activation time constants *τ*_*E*_ and *τ*_on_, whereas coupling strengths *k*_HAPC_ and *k*_CMP_ produced negligible changes in these global metrics near the healthy operating point. This does not imply that coupling is unimportant: in regimes such as STC, reduced *k*_*κ*_ disrupts phase coherence and wave coordination even if peak pressure and motility index remain locally insensitive around the healthy baseline. In contrast, the duty fraction *ϕ*_duty_ and distal gating parameters (e.g., *α*_spread,CMP_, *p*_on_, *p*_off_, *τ*_*P*_) showed relatively modest effects. Overall, these findings emphasize the dominant role of proximal excitation in shaping whole-colon pressure patterns. Because quantitative metrics of spatiotemporal organization are difficult to define, sensitivity was evaluated using only global pressure measures.

**Fig. 3.**
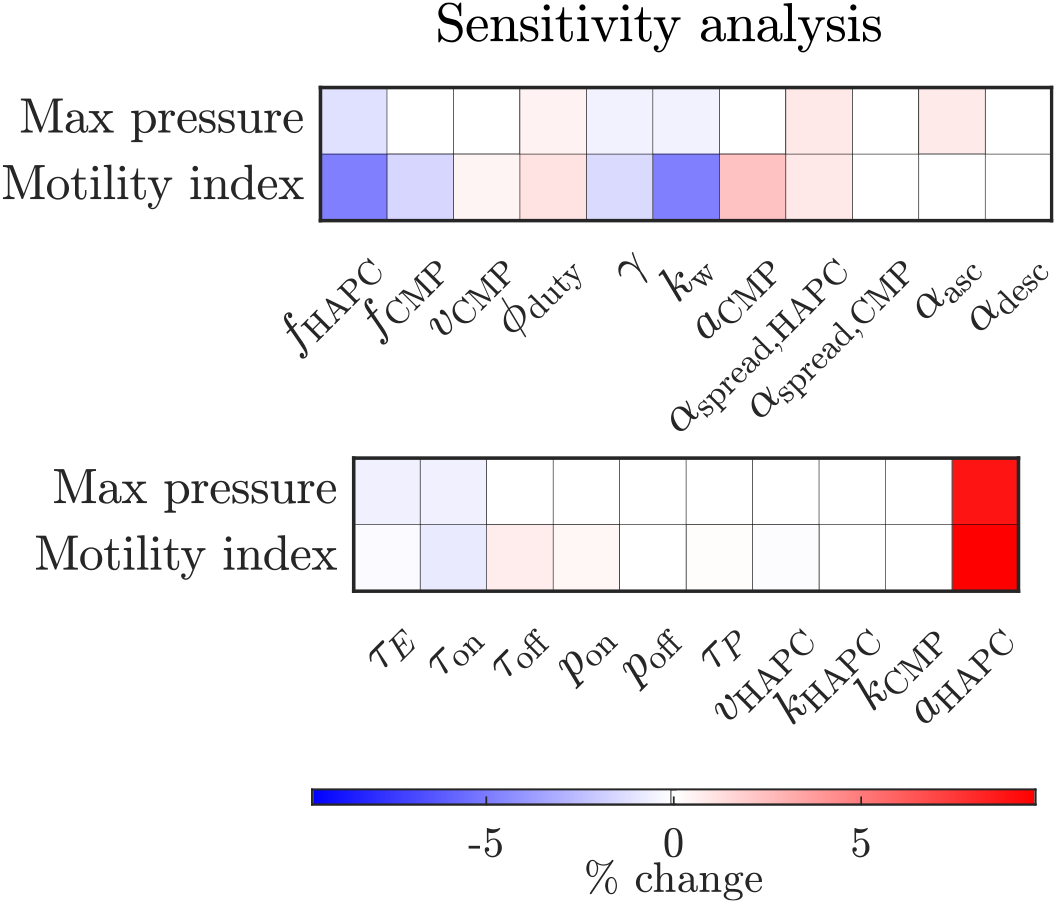
Sensitivity analysis heatmap Each parameter was independently increased by 10% relative to the healthy baseline, and changes in peak pressure and motility index are shown Sensitivities represent local responses of the nonlinear model and should be interpreted as relative effects near the healthy regime rather than global trends

## 4 Discussion

IBS and chronic constipation are common functional gastrointestinal disorders that contribute substantially to morbidity and healthcare burden [1–4, 6]. High-resolution colonic manometry has revealed rich spatiotemporal motor patterning in humans and has clarified how these patterns are altered in disease [7, 8]. Yet, because manometry reflects the combined output of underlying pacemaker activity, enteric reflexes, and colon wall mechanics, the mechanisms that generate and disrupt these patterns remain difficult to isolate in vivo [48]. To bridge this gap, we developed a biophysical model that integrates key elements of colonic physiology, including interstitial cell pacemakers, enteric reflexes, and rectosigmoid braking, to reproduce motility patterns observed in vivo. The model offers a mechanistic framework for linking manometric data to underlying control processes, enabling both hypothesis generation and the interpretation of disease-specific abnormalities. The correspondence between simulated and clinical pressure maps in Fig. 2 illustrates the validity of the framework.

### Model novelty

Our whole–colon model reproduces illustrative examples of clinically observed healthy, IBS–D, and STC-like motility patterns by tuning a subset of parameters: wave amplitude and propagation speed, inter-segmental coupling, stochastic phase noise, and distal gating. Prior models have captured important aspects of this system; for example, weakly coupled oscillator frameworks establish how ICC network coupling and frequency shape coordination [28, 29], and neurogenic/peristaltic models emphasize ascending excitation and descending inhibition [26, 30]. However, these models often focus on short segments, lacking explicit ENS-ICC-smooth muscle integration, or they utilize animal systems that limit human translation. Our approach incorporates (i) regional specialization (proximal mixing vs distal storage/emptying) [7, 49], (ii) reflex asymmetry (oral excitation and anal inhibition), also known as the law of the intestine [37], and (iii) a pressure–gated rectosigmoid modulation element supported by modern high-resolution manometry[13, 16]. To our knowledge, this is the first model that reproduces illustrative examples of multiple human motility patterns (healthy, IBS–D, and STC) through physiologically interpretable parameter changes.

### Healthy case

The model reproduced key motility patterns observed in healthy colonic function: intermittent HAPCs traversing long distances and retrograde CMPs in the distal colon that implement rectosigmoid brake dynamics. In vivo high-resolution manometry studies show that HAPCs are large amplitude, long–range events often triggered by waking or meals that mediate bulk transit through the colon and aid in defecation [7, 10, 32, 50]. The model reproduces HAPCs with frequencies and amplitudes consistent with those reports. In the distal colon, the model generated retrograde CMPs that oppose the antegrade movement of luminal contents into the rectum; such patterns have been identified by high-resolution manometry as a key component of the rectosigmoid brake and continence mechanisms [13, 16]. The overall pattern of anal propulsion in the proximal colon and regulated braking in the distal colon reflects the spatial organization and functional roles observed in healthy humans [7, 8].

### IBS-D case

In the illustrative IBS-D simulation, increasing the wave speed and reducing distal gating produced a hypermotile, poorly regulated pattern. A distal pressurization emerged when upstream contractions encountered segments that failed to relax. Although the simulated traces do not display a focal pressure spike (as pressure is determined locally by wall mechanics without explicit mass transport), the nonlinear tube law still helps explain how advancing contents into a stiffened distal segment can disproportionately elevate pressure. In addition to the reduced luminal area, local increases in wall stiffness due to muscular contraction or incompressible content further amplify the pressure response. This mechanism could explain clinical observations: in IBS-D patients, impaired postprandial coordination of retrograde rectosigmoid activity may lead to ineffective clearance of intraluminal gas, resulting in its accumulation and entrapment [16]. When combined with visceral hypersensitivity, this localized gas retention may provoke postprandial discomfort, pain, and bloating [51]. Clinically, IBS patients frequently report pain at lower distension thresholds than healthy individuals, and heightened contractile and sensory responses are well documented [52]. Our model reproduces these hallmark features and provides a mechanistic explanation that links abnormal motility coordination with symptom generation in IBS-D.

### STC case

In the illustrative STC case, reducing coupling and introducing phase noise disrupted pacemaker synchrony and abolished long-range coordination, resulting in low-amplitude, non-propagating contractions with markedly diminished HAPCs features consistent with manometric profiles in STC [25, 45]. Histological and modeling studies implicate ICC depletion and, thereby, impaired network coherence as possible underlying factors in this response [28, 29]. Our abstraction captures this through a simple mechanism: fragmented pacemaker activity (phase noise) combined with weak coupling prevents wave superposition and coordinated propagation. The outcome is sparse or absent HAPCs, a low motility index, and delayed transit, which closely match the clinical findings in STC patients [25, 45].

### Generalizability of our model

A key strength of our model is its generalizability: by adjusting a few interpretable parameters (ICC drive/coupling, neural reflex, distal brake, and phase noise), the same model generates multiple clinically observable motor patterns. Sensitivity analysis shows that parameters governing proximal drive and waveform morphology dominate global pressure metrics near the healthy baseline, whereas distal gating and reflex gain play a more modest role in this regime. This supports the view that diverse patterns could potentially emerge from modest shifts in a shared control architecture. However, it is important to note that the influence of some parameters (e.g., HAPC pacing frequency) depends on their interaction with waveform shape and activation dynamics, emphasizing that sensitivities are local to the operating point rather than globally monotonic. As such, our model is a hypothesis generation tool; for example, strengthening distal inhibition should mitigate IBS–D pressure spikes; enhancing ICC coupling should improve STC propagation. Both propositions follow directly from the model and are testable experimentally (e.g., via neuromodulation or pharmacology) [12, 53].

### Future directions

#### Patient-specific calibration

One potential application of the model is to fit parameters to individual high-resolution manometry recordings, such as coupling strength, reflex gains, and baseline tone, to generate virtual, patient-specific colon models. This approach could help infer the mechanistic contributions of various deficits, such as impaired ICC or ENS function [7, 53].

#### Modeling additional motility patterns

The current model can be extended to explore other clinically relevant motility patterns, such as opioid-induced constipation [54] and post-surgical dysmotility [55]. By modifying regional parameters in physiologically plausible ways, the model could simulate whether the observed motility disruptions are explainable within the existing framework.

#### 3D/FSI mechanics

Future versions of the model could incorporate three-dimensional continuum mechanics or fluid–structure interaction (FSI) to simulate pressure–flow coupling, wall stress, and mixing. Recent gastrointestinal electromechanics and FSI studies highlight the utility of such approaches in linking mechanical forces to function [26, 27].

#### ENS and mechanosensing

The model could be refined by incorporating more detailed representations of the enteric nervous system, including interneuronal circuits [56] and mechanosensory pathways [57]. These additions would further enhance fidelity and support a more comprehensive multiscale ICC–ENS–muscle framework [33, 53].

### Limitations

The model uses simplified oscillators and a one-dimensional discretization to approximate the spatial and electrical complexity of real ICC and enteric neural networks [28, 29]. The distal ‘rectosigmoid brake’ component is implemented as a phenomenological abstraction inspired by distal motor patterns observed in high-resolution manometry; the precise physiological circuitry underlying this braking behavior remains incompletely defined. The division into proximal and distal subnetworks represents a functional approximation of regional specialization. In vivo, however, transitions are gradual and mediated by distributed enteric neural circuits, as demonstrated in mice [58, 59], and the equivalent circuit-level organization in the human colon remains an active area of investigation. Accordingly, although the parameter sets reproduce clinically recognizable manometric patterns, they were chosen to be physiologically plausible and to match the qualitative features of the target pressure maps rather than being uniquely inferred from data. Because several parameters (e.g., effective ICC coupling *k*_*κ*_, spread *α*_spread,*κ*_, and gating thresholds *p*_on_, *p*_off_) are difficult to measure directly, and because multiple parameter combinations can generate similar pressure patterns, the reported values should be interpreted as example ranges rather than definitive physiological measurements, consistent with the approach in other reported motility models [60]. In addition, propagation speed is prescribed through the per-segment phase lag δ_*κ*_ (and the corresponding initial phase gradient) rather than emerging entirely from oscillator dynamics.

Given that several intermediate states governing wave selection are not directly observable, we adopt a hybrid mechanistic–phenomenological framework that captures the major clinically observed motor patterns while remaining physiologically interpretable. It is important to note that the simulated cases represent only three illustrative examples of clinically observed motility patterns. In practice, patients exhibit substantial physiological variability, and no single parameter set can capture the full spectrum of clinically observed behaviors. Systematically exploring this broader range of parameter combinations is a substantial task and is left for future work.

We developed a simple, biophysically based model of colonic motility that reproduces illustrative examples of healthy function, as well as IBS-D-like and STC-like motor patterns. Through physiologically reasonable parameter adjustments, the model generates pressure patterns that align with clinical high-resolution manometry in terms of spatial organization, propagation, and amplitude. It also offers mechanistic explanations for symptom generation; for example, how impaired rectosigmoid braking and nonlinear wall mechanics may lead to distal pressurization in IBS-D. By accounting for multiple illustrative patterns within a single framework, the model supports the view that motility disorders could potentially arise from shifts in shared physiological mechanisms and provides a foundation for hypothesis testing, mechanistic studies, and patient-specific applications.

## Acknowledgements

A.A.K acknowledges email communications with Dr. Sean Parsons (McMaster University) regarding ICC networks and technical discussions with Dr. Daniele Schiavazzi (University of Notre Dame) on modeling fluid–structure interaction, Dr. Martina Bukač (University of Notre Dame) on tube law, and Dr. Robert Rosenbaum (University of Notre Dame) on modeling neural circuits.

## Statements and Declarations

### Supplementary information

No supplementary material.

### Ethics approval and consent to participate

This study reused de-identified high-resolution manometry recordings that were collected as part of our previous studies, for which institutional ethics approval and informed consent were obtained. No new ethics approval or consent was required.

### Consent for publication

Not applicable

### Data availability

Not applicable

### Materials availability

Not applicable

### Code availability

The model developed in this paper is available at this github link: https://github.com/ahilanananthakrishnan/ABEManometry

### Author contribution

Conceptualization: A.A.K., P.G.D.; Methodology: A.A.K.; Formal Analysis: A.A.K.; Investigation: P.G.D.; Resources: M.A.H., P.G.D.; Writing – Original Draft: A.A.K.; Writing – Review & Editing: A.A.K., P.G.D., M.A.H.; Supervision: P.G.D., M.A.H.; Project Administration: M.A.H.; Funding Acquisition: M.A.H.

